# Early vegetation recovery after the 2008-2009 explosive eruption of the Chaitén Volcano

**DOI:** 10.1101/746859

**Authors:** Ricardo Moreno-González, Iván A. Díaz, Duncan A. Christie, Rafael E. Coopman, Antonio Lara

## Abstract

In May 2008, Chaitén volcano entered in eruptive process, one of the world largest eruptions in the last decades. The catastrophic event left different type of disturbance and caused diverse environmental damage distributed heterogeneously in the surrounding areas of the volcano. We went to the field to assess the early vegetation responses a year after the eruption, in September 2009. Particularly, we evaluated the lateral-blast disturbance zone. We distributed a set of plots in three disturbed sites, and one in an undisturbed site. In each of these sites, in a plot of 1000m^2^ we marked all stand tree, recording whether they were alive, resprouting or dead. Additionally, in each site 80 small-plots (~4m^2^) we tallied the plants regeneration, its coverage, and the log-volume. We described whether the plant regeneration was growing on mineral or organic substrate. In the blast-zone the eruption created a gradient of disturbance. Close to the crater we found high devastation marked by no surviving species, scarce standing-dead trees and logs, as well as no tree regeneration. On the other extreme of the disturbance gradient, the trees with damaged crown were resprouting, small-plants were regrowing and seedlings were more dispersed. The main regeneration strategy was the resprouting of trunks or buried roots, while few seedlings were observed in the small plots and elsewhere in disturbed areas. However, the assessment was too soon after the eruption and updated monitoring is required to confirm observed patterns. The main findings of this study are: i) a mosaic of pioneering-wind dispersed species, scattered survivors regrowing and spreading from biological legacies, and plant species dispersed by frugivorous birds, likely favored by the biological legacies; (ii) the early succession is influenced by the interaction of the species-specific life history, altitudinal gradient and the different intensity of disturbance.

## 1. Introduction

Volcanic eruptions are large-scale disturbance that modify the composition, the structure and ecosystems processes (Swanson et al., 2013; Turner and Dale, 1998; Turner et al., 1997). As volcanoes enter in eruptive phase complex mechanism involve the events in a gradient of multiple type of disturbances such as explosive blasts, thermal and toxic chemical waves, landslides, glowing avalanche deposits, debris flows, lava flows, and air falls of volcanic tephra (Dale et al., 2005; Swanson et al., 2013). A volcanic eruption could have effects at very large spatial and temporal scales (Peet, 1992; Robock and Oppenheimer, 2003), yet directly on surrounded forest throughout burning, burying and/or blowing down trees or other plants (Dale et al., 2005). However, since this kind of large disturbance are infrequent (Turner et al. 1998) or have been seldom monitored (Chrisafully and Dale, 2018), our understanding of vegetation responses is still limited. Despite the importance of volcanic eruptions, few studies have analyzed the effect of eruptions short after the event (Dale et al., 2005; Chrisafully and Dale 2018). On the other hand, the effect of volcanism has been studied by comparing different stands in chronosequences assuming that different forest patches would reflect the successional stage after the disturbance (e.g. Drury and Nisbet 1973; Franklin and Swanson, 2010; Pickett 1988; Walker et al 2010).

Forest recovery is considerably slow following disturbances that heavily impact soils as well as aboveground vegetation (Chazdon, 2003). Despite volcanoes are considered as catastrophic disturbance event, it is possible that diverse biological legacies can remain after the event. The persistence and heterogeneous distribution of these biological legacies might modulate and help the recolonization of the plants (Foster et al., 1998; Franklin et al., 1985; Turner et al., 1998). For example, the life-history traits of the pre-disturbed community will result in particular survivors perhaps able to re-colonize and expand over the new conditions on the disturbed environment (Foster et al., 1998; Turner et al., 1998). Therefore, the arrange and the amount of the biological legacies would drive the rate and the pathways of the new colonizers.

One of the most active tectonic and volcanic areas in the world correspond to the subduction area of the Nazca and the South American plate (e.g., Stern, 2004), which also originated the longest and the second highest mountain range in the world, the Andes. The volcanoes in the Chilean Andes are quite active. The Southern and Austral Volcanic Zones comprise 74 active volcanoes since the Post-Glacial up to the present and at least 21 have had one large explosive eruption (Fontijn et al., 2014). In the southern portion of the Andes, it is located the South American Temperate rainforest (SATR) (Armesto et al., 1998). This cordillera plays a critical role in SATR dynamics, where current successional models after volcanic disturbance suggest 80 the Nothofagus are dominant pioneer trees that benefit from disturbances to persist (Veblen et al., 2016). The SATR represent a biogeographical island, rich in endemic species and dominated by evergreen, broad-leaved species (Armesto et al., 1996; Villagran and Hinojosa, 1997, Arroyo et al., 2004). South of 40°S, extensive areas of old-growth forest dominate the mountainous landscape. These forest are not directly disturbed by modern human activity, and represent one of the last pristine areas with pre-industrial biogeochemical conditions (Perakis and Hedin 2002)

On May 1^st^ 2008 started the last eruption of the Chaiten Volcano (42° 50’S), near the small town of Chaiten, in southern Chile. This eruption was the largest rhyolite eruption since the great eruption of Katmai Volcano in 1912, being the first rhyolite eruption in which some attributes of its dynamics and impacts have been monitored (Major and Lara, 2013). The eruption consisted of an approximately 2-week-long explosive phase that generated as much as 1 km^3^ bulk volume tephra (∼0.3 km^3^ dense rock equivalent) followed by an approximately 20-month-long effusive phase that erupted about 0.8 km^3^ of high-silica rhyolite lava that formed a new dome within the volcano’s caldera (Major and Lara, 2013). Chaiten volcano was surrounded by extensive old-growth undisturbed forest by humans, providing a unique opportunity to evaluate the effects of volcanism over undisturbed landscape, then providing an opportunity to understand how volcanism may have affected the forest, under pre-industrial conditions.

In this study, we characterized the early establishment of the vegetation after the eruption of the Chaitén volcano along a disturbance gradient one year after the eruption started, from nearby the crater to the closest old-growth forest with vegetation completely alive. We predict that plant species diversity and the number of living trees decrease close to the crater, and pioneering *Nothofagus* species are colonizing the areas affected by the eruption. Our objectives included the creation of a baseline for further long-term monitoring of vegetation recovering, and to identify and analyze the importance of biological legacies for forest recovery.

## 2. Material and methods

### 2.1 Study site

The Chaitén volcano is located in the west side of the Andean Range in southern Chile, at 42° 50’ S and 72° 39’ W (Fig 1) and is part of the so-called Southern Volcanic Zones. The Chaitén Volcano is a dome of 1100 m height, located at 10 km NW of the town of Chaitén, populated by approximately 5000 inhabitants before the eruption. The climate of the area corresponds to a humid temperate with a strong oceanic influence (Luebert and Pliscoff, 2006). The landscape is characterized by sharp mountains shaped by glaciers and valleys originated by fluvial and glacio-fluvial outwash (Swanson et al., 2013). Vegetation within the influence area of the Chaiten volcano is dominated by a dense old-growth temperate rainforest of northern Patagonian and Valdivian types (Luebert and Pliscoff, 2006; Swanson et al., 2013; Veblen et al., 1983; Veblen et al., 1996). These forests composition is dominated by broad-leaved evergreen trees such as *Nothofagus dombeyi* (Mirb.) Oerst. (Nothofagaceae), *Laureliopsis philippiana* (Looser) Schodde, *Gevuina avellana* (Molina), *Amomyrtus luma* (Molina) D. Legrand and Kausel, *Luma apiculata* (DC.) Burret, *Drymis winteri* J.R. Forst. and G. Forst., *Eucryphia cordifolia* Cav. and *Weinmannia trichosperma* Cav. (both Cunoniaceae). These species are normally arranged in multi-layered canopy strata and old, large-emergent trees (up to c. 40 m height) and abundant snags and logs. The understory is densely covered by ferns such as *Lophosoria quadripinnata* (J.F.Gmel.) C.Chr. and *Blechnum magellanicum* (Desv.) Mett., seedlings, saplings and bamboo tickets (*Chusquea* spp.).

**Figure 1.**
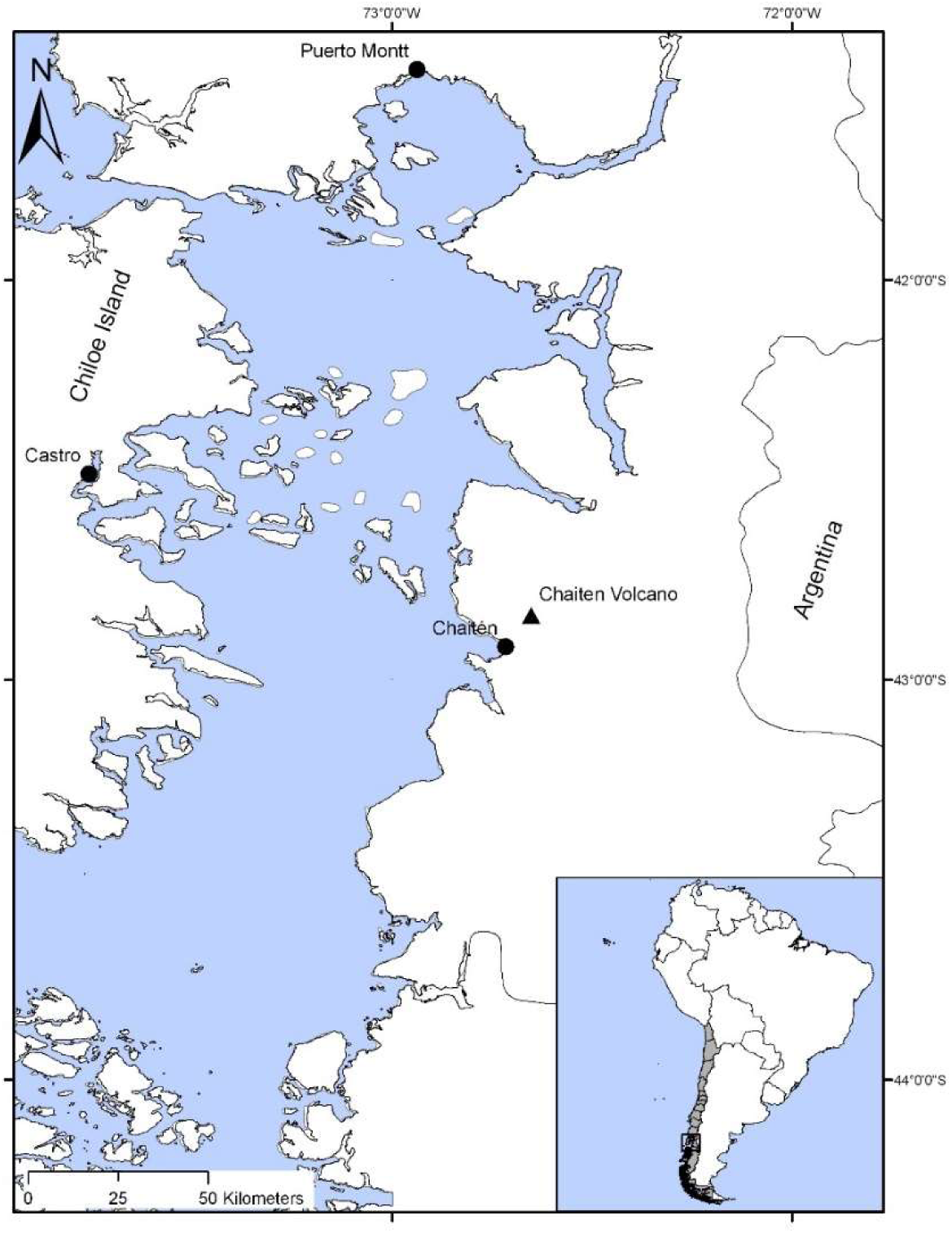
Study area.

### 2.2 Study design

We carried out the field work in September 2009. We first conducted a general inspection of the study area (del Moral, 1981). Later, we selected four sites with different degrees of volcanic impact following similar classifications of previous studies (del Moral, 1981). The first site corresponds to a forest with low volcanic impact that only received few millimeters of tephra deposition. This site was called “Old-growth forest”. It was located 5 km from the center of the new dome at an elevation of 600 m a.s.l. (Figure 1b). The second site was characterized by the presence of abundant dead standing trees, and higher tephra deposition. This site was called “Standing-dead” located at 3.5 km from the center of the new dome, at 160 m. The third site was more affected by the eruption, and was characterized by high tephra deposition, where most trees were broken and throw down by the eruption. This site was called “Blow down” and was located at 2.3 km from the center of the new dome at an elevation of 300 m a.s.l. Finally, the fourth and most affected site was in the border of the crater, at 1.5 km of the center of the new dome and at 750 m a.s.l. This site presented the higher tephra deposition and all trees were blow down by the eruption and heavily buried under volcanic ash. This site was called “Total destruction”.

In each one of these four areas was installed a 50 × 20 m permanent plot where we measured the diameter at the breast height (dbh) of all trees ≥ 5cm. Every tree was identified at the species level when possible and marked. We registered if the stems were alive, resprouting after the eruption, or snag (standing dead). To estimate the presence and abundance of tree regeneration, we traced four transects of 120 m long. Transects were distributed parallel, separated 20 m from each other. In each transect, we mounted five circular plots of 1.5m radius (4.71m^2^) every 30 m. On these plots we counted all seedlings and saplings present, and recording the substrate where they rooted (forest soil, logs, or volcanic tephra). We registered other non-tree plant species in the circular plots, estimated its abundance as the relative cover, and we also recorded the substrate where they rooted were organic or mineral (e.g., forest soil, logs, or volcanic tephra). We assessed the volume of all exposed logs in the plot, until each border within the plot. Assuming a cylindrical form of each log, we calculated the volume of logs present inside the plot and standardizing its measurements to m^3^ of logs per m^2^ of plot surface.

### 2.3 Data Analysis

Trees seedlings, herbs and shrubs species richness was compared among sites using rarefaction analysis (Colwell 2006). Rarefaction curves allow for comparing species richness among environments avoiding the bias in the number of species detected due to the increased probability of species detection in areas where individuals were more abundant, by plotting the number of species detected in function of the number of individuals observed (Colwell and Coddington, 1994, Gotelli & Colwell 2011, Colwell et al., 2012). This procedure includes Monte Carlo simulations delivering an average value of species richness with a confidence interval of 95% (Colwell 2006). Rarefaction analyses were conducted using the software EstimateS Win 8 (Colwell 2006). Non-vascular species were excluded from the rarefaction analyses since not all of them were classified up to the species level, and one genus could include several species. Plant composition among sites was compared using Sørensen’s similarity index based on the presence/absence of plant species. Rarefaction and similitude analyses were both carried out with the software Estimates 8.2 (Colwell 2006). In attempting to define if the plant composition of one site was a subset of the composition of other site, we conducted a Nested Analysis in NestCalc free access software (Atmar and Patterson, 1995).

## 3. Results

### 3.1 General description of the effects of the eruption

The eruption of the Chaitén volcano has an impressive effect on the forest. The eruption in the study area produced a gradient of destruction. Around the crater all the trees were blown down and heavily buried by tephra. Other trees were partially buried, broken in half, with the trunks covered with impacts of many little stones looking similar to the fire of gun shots (Fig x). Moving away and down from the crater, trees were blown down and densely covered by ash(Fig x), while further away trees were standing dead presenting progressively finer branches(Fig x). Many small patches of the organic litter were exposed and sparsely distributed and protected from the direct effect of the eruption by root discs and large trunks, or were exposed by small ravines that flush away the volcanic ash. These small patches looked like clumps of organic soil, with roots of ferns and other small plants that were not buried by the tephra. These clumps were covered by plants that survived in the area affected by the eruption, and their frequency increases when moving away from the crater towards the undisturbed forests.

In the surrounding, we did not observed evidence of fire caused by the eruption, but the trunk and bark of several trees looked charred within the influence area. Trees were dead and dry by the eruptive explosions, but some trees in Blow-down and Standing-dead areas maintained the bark in the opposite side of the crater, perhaps with possibilities to resprouting hereafter. Tephra buried the fallen trees in the whole study area, but principally in Blow-down and Total-destruction areas. Tephra was very compacted, seeming as pavement that covered the fallen forests. Heterogeneous tephra deposition exposed some trunks, clumps and remnants with organic-forest soil likely from the original forest. Shrubs and epiphytes were spread everywhere in the Standing-dead area. Ferns, bamboo and tree seedlings were growing in remnants of exposed organic soil, usually below or at one side of trunks or unearth roots. Surviving trees also increases when moving away and down from the crater.

### 3.2 Species composition

We found 10 species of trees in the area, and 50 species of smaller plants, including seedlings, herbs and ferns (Table 1). Rarefaction analysis for tree species showed that the Standing-dead plot holds one more species than the Old-growth forest plot (Fig. 3). This same analysis showed the Blow-down plot holds very few individuals, but the initial slope and shape of the curve was similar to the other two curves. It indicates that at similar number of individuals, all curves showed a similar number of tree species. Rarefaction analysis of small vascular plants showed the Old-growth forest site had the richest species composition, while decreasing in the Standing dead and Blow-down plots (Fig. 3). The Total-destruction plot showed practically no species.

**Table 1.**
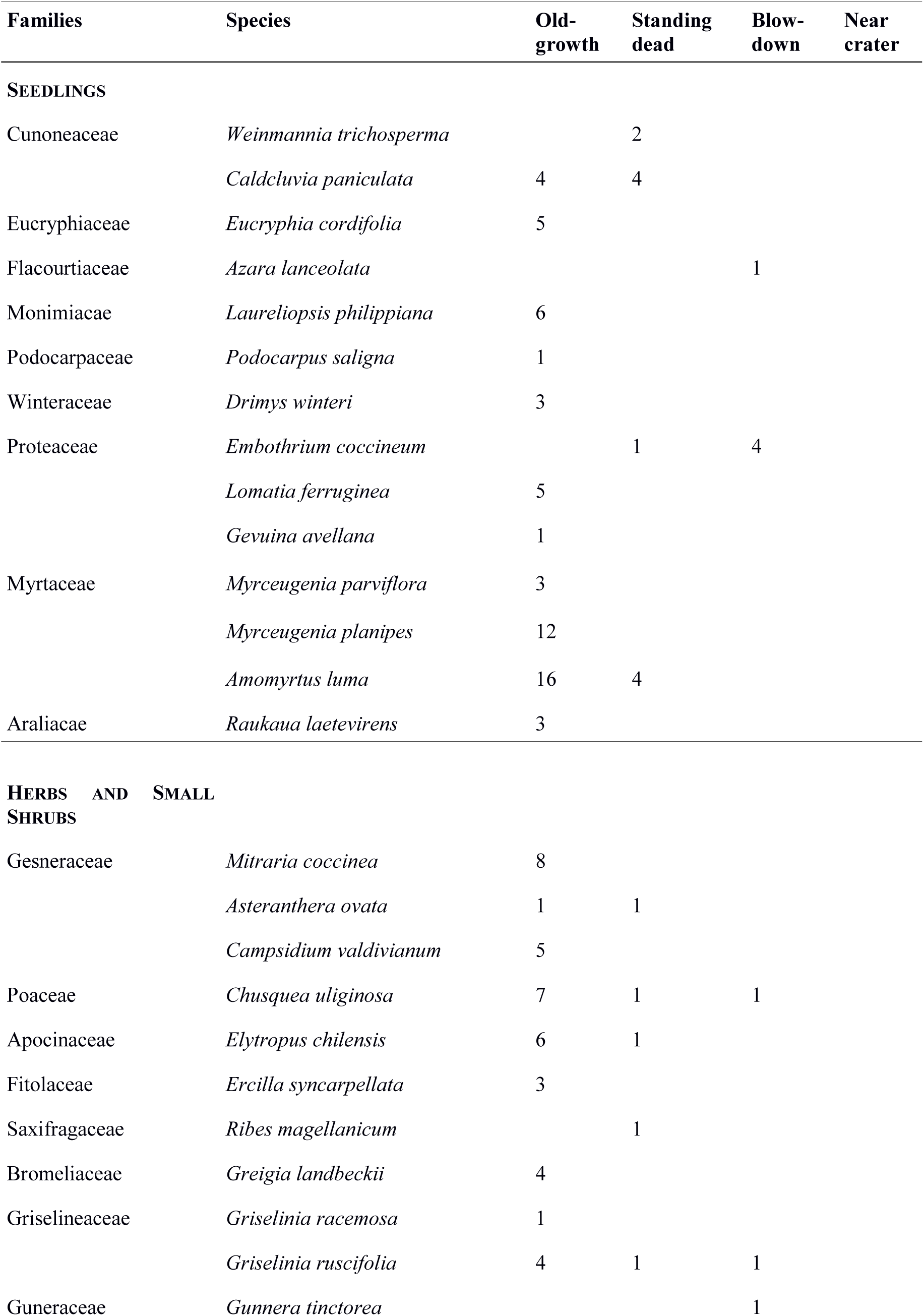
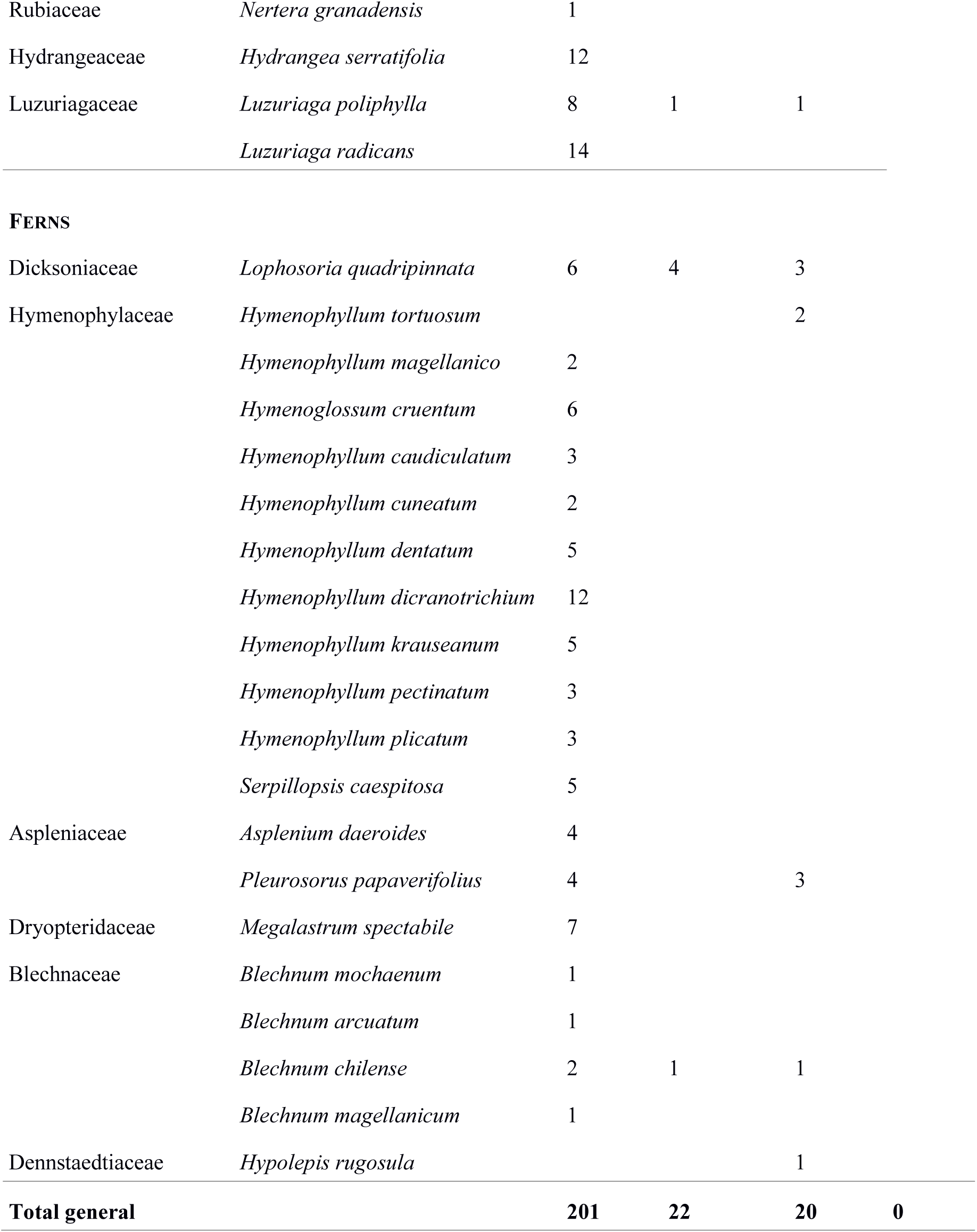
List of plant species present in the Chaitén volcano and their frequency in the big plot and in 20 plots of 3.14m^2^ in the different studied areas.

**Figure 2.**
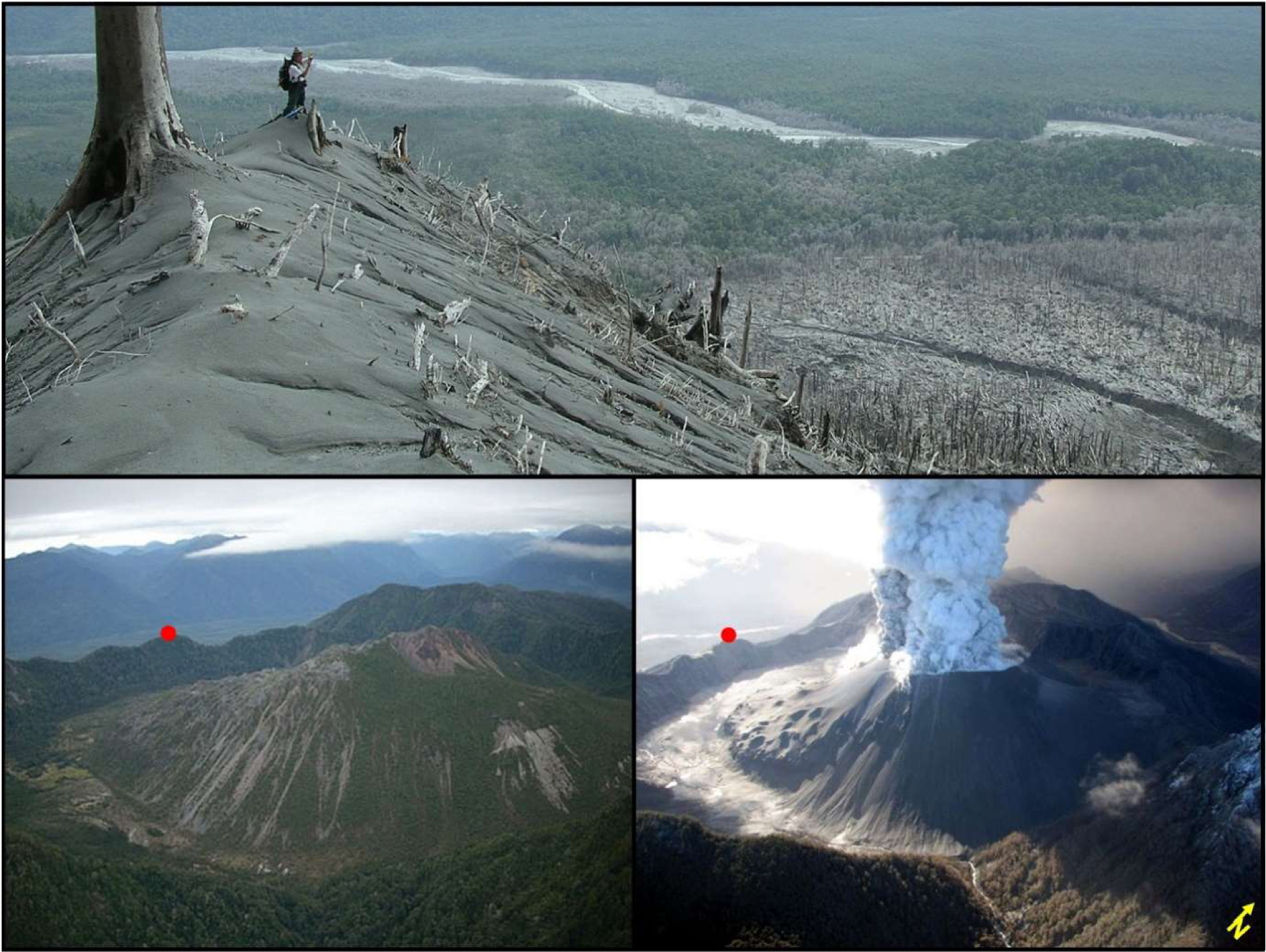
Upper panel showing impressive impacts close to the crater. Comparison of the volcanic dome and vegetation cover before (bottom-left panel) and during the 2008-2009 eruption (bottom-right panel).

**Figure 3.**
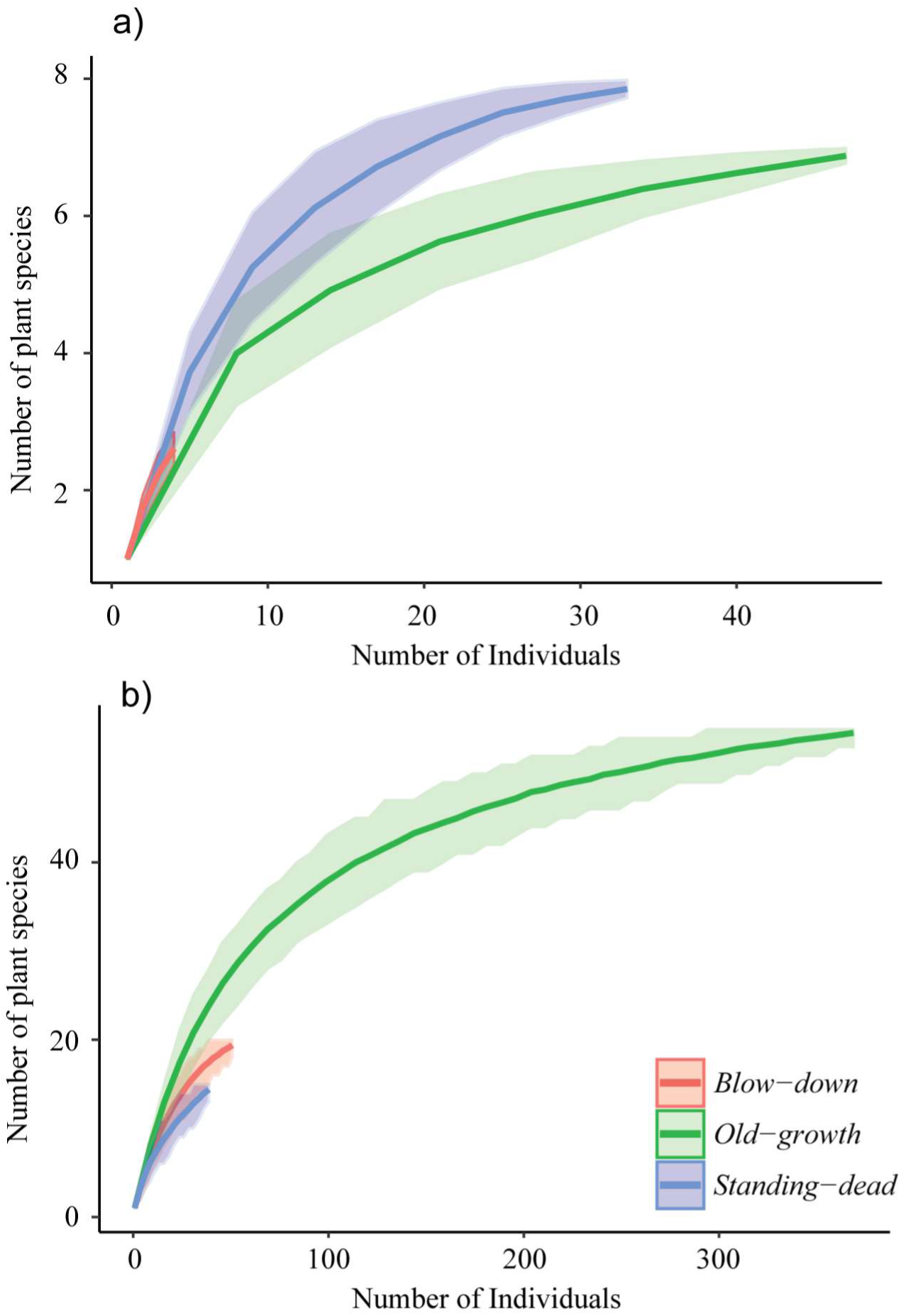
Rarefaction analysis showing the relationship between number of tree seedlings species (a) and small vascular plant species (b) as function of the number of individuals. Both panel the shaded area is indicating one standard error). All sites affected by the eruption of Chaitén Old-growth, Standing-dead and Blow-down are compared, except Total-destruction was not included because of the absence of plants.

Plant composition among Old-growth and Standing-dead plots showed some similarities (34%), but less related with Blow-down (20%) and the Total-destruction plot (0%). Standing-dead and Blow-down plots showed the highest similarities in species composition (Table 2). In the disturbed plots, the plant species recorded were a subgroup of the species present in the Old-growth forest plot (Nestedness calculator T= 13.85°). The most frequent plant was the fern *L. quadripinnata* that survived the direct impact of the eruption in small clumps of organic matter protected by trunks or root disc of trees, and were not completely covered by tephra. Bamboo and ferns were sprouting from roots in the remnants of exposed forest soil.

**Table 2.**
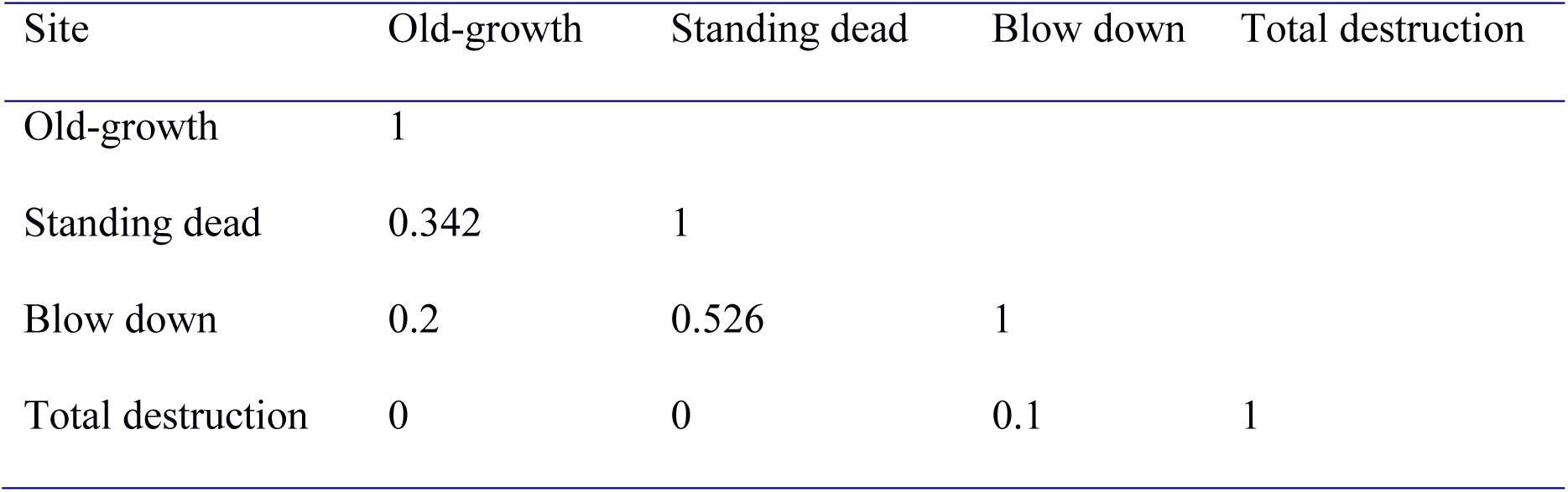
Sørensen similarity index for the plant composition among the different sites in the Chaitén influence area.

### 3.3. Forest structure

The forest structure of the Old-growth site showed individuals of all sizes, with several large emergent trees (Fig. 4). Most trees in the Old-growth plot were alive and the snags most probably represent trees that died before the eruption, while in the other plots most trees were killed by the eruption. The Standing-dead plot showed similar dbh distribution than the Old-growth plot, with higher frequency of small dbh. The Blow-down plot showed a very small number of standing trees, concentrated in intermediate size class, while the Total-destruction plot practically had no individual standing trees (Fig. 4).

**Figure 4.**
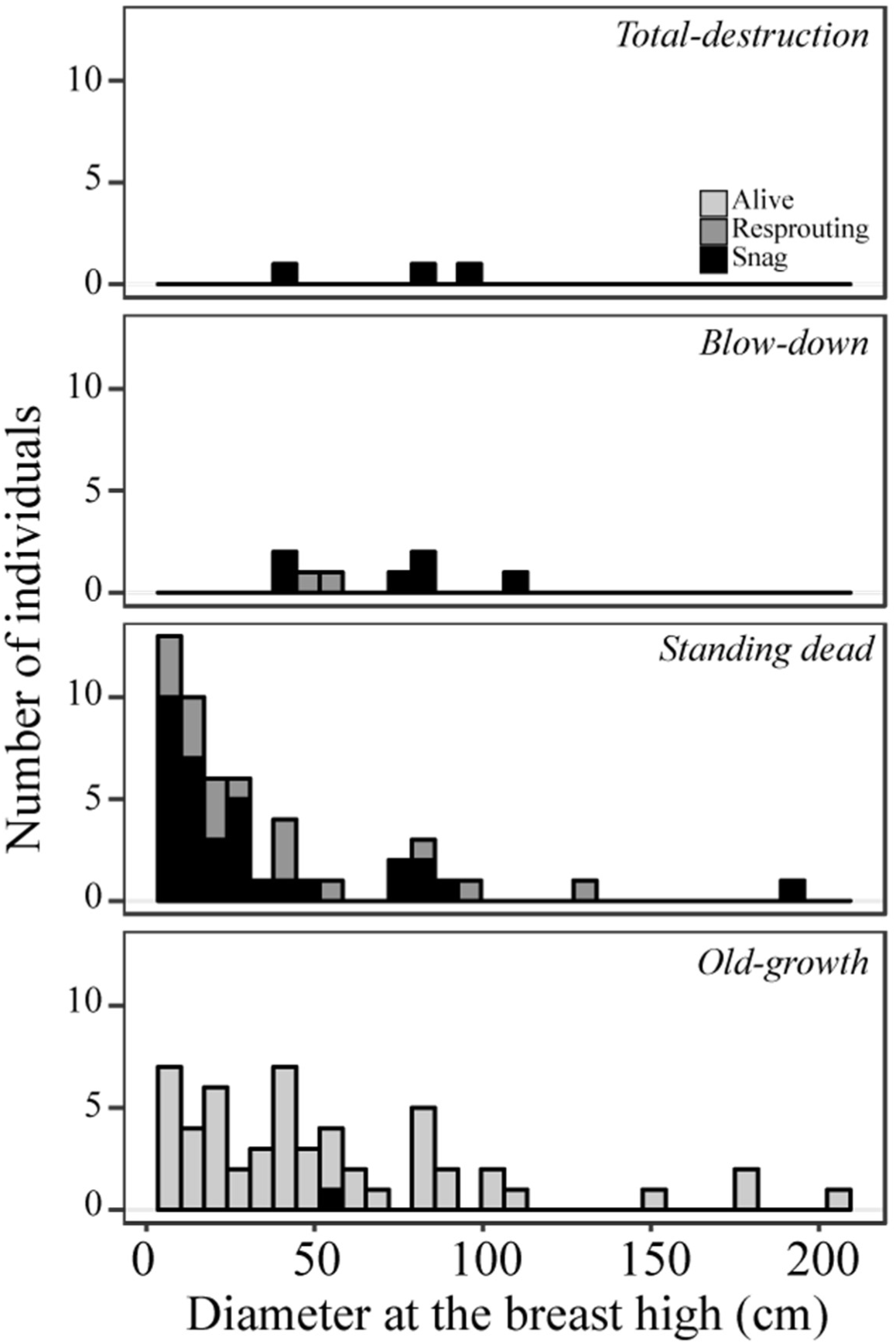
Diameter at the breast height (dbh) distribution for living and standing dead trees in the study sites influenced by the eruption of the Chaitén volcano.

The basal area of the Old-growth plot was dominated by *L. philippiana*, followed by *E. cordifolia* and *C. paniculata* (Table 3). In the Standing-dead and Blow-down plots *W. trichosperma* and *N. dombeyi* were the dominant components of the basal area, but also with a high number of unidentified trees (Table 3). We were not able to identify the species of the standing dead trees in the Total destruction plot. The volume of fallen trees increased in the Blow-down and in Standing-dead plots, but it was much lower in the Old-growth and the Total-destruction plots (Fig. 5). In the Total-destruction plot most logs were heavily buried by a dense tephra layer.

**Table 3.** Total basal area in 1000 m^2^ of tree species of the study plots in the Chaitén volcano area, classified as alive, resprouting after the eruption, or snag

|  | Total-destruction | Blow-down |  | Standing-dead |  | Old-growth |  |
| --- | --- | --- | --- | --- | --- | --- | --- |
|  | Snag | Snag | Resprouting | Snag | Resprouting | Snag | Alive |
| *Amomyrtus luma* |  |  |  | 0,02 | 0,34 |  | 0,47 |
| *Amomyrtus meli* |  |  |  |  | 0,06 |  |  |
| *Caldcluvia paniculata* |  |  |  |  |  |  | 2,54 |
| *Drimys winterii* |  |  |  | 0,14 |  |  |  |
| *Lomatia ferruginea* |  |  |  | 0,,04 | 0,01 |  | 0,02 |
| *Myrceugenia planipes* |  |  |  |  |  |  | 0,84 |
| *Eucryphia cordifolia* |  | 0,55 |  | 0,59 | 0,61 | 0,26 | 6,37 |
| *Laureliopsis philippiana* |  | 0,25 |  | 0,87 |  |  | 9,64 |
| *Nothofagus dombeyi* |  |  |  | 3,46 |  |  |  |
| *Weinmannia trichosperma* |  | 0,95 | 0,41 |  | 2,38 |  | 0,71 |
| *not identified* | 1,41 | 0,94 |  | 1,45 |  |  |  |
| **Total** | **1,41** | **2,69** | **0,41** | **6,57** | **3,39** | **0,26** | **20,59** |

**Figure 5.**
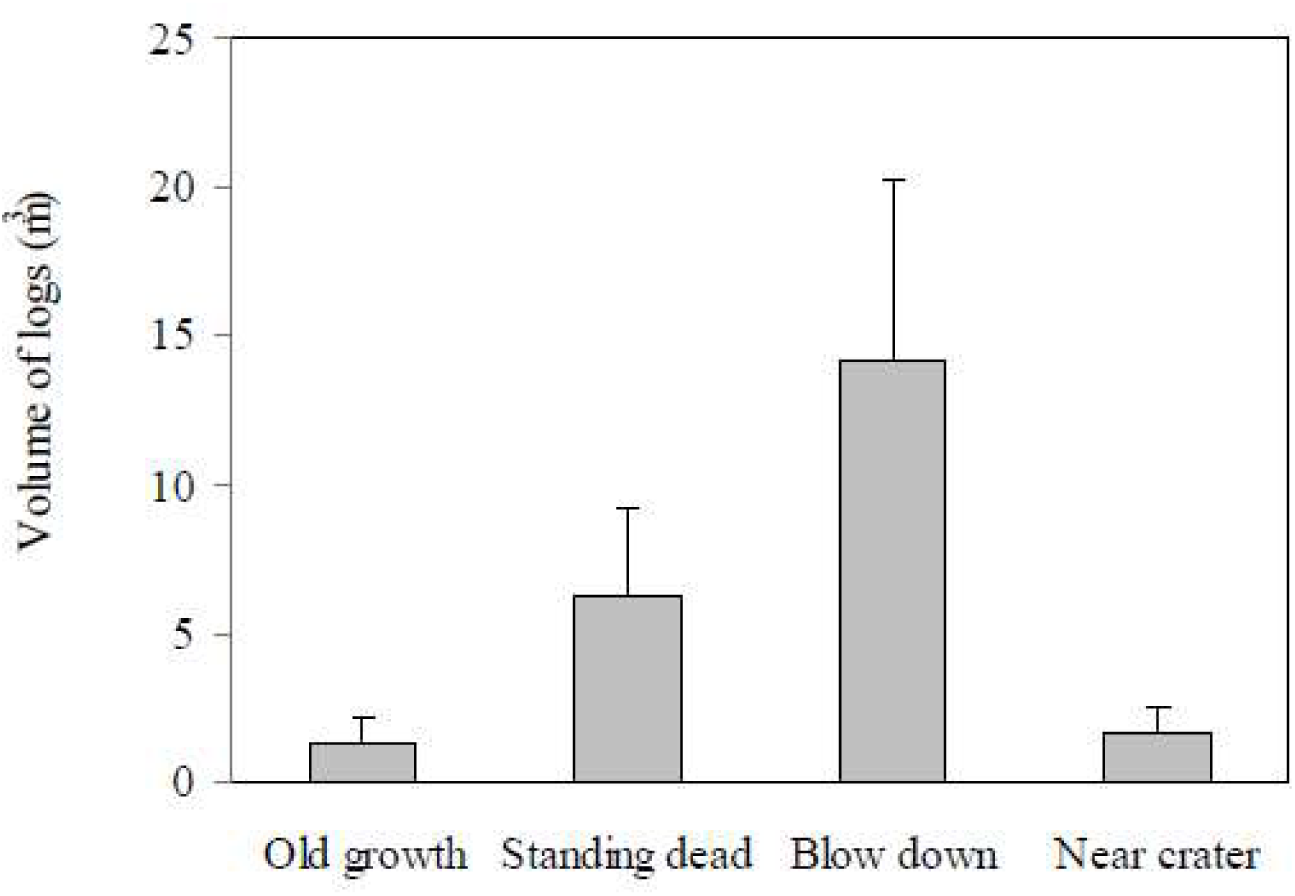
Volume of fallen dead trees (logs) in the study sites influenced by the eruption of the Chaitén volcano in the study sites.

### 3.4. Seedlings and small plants

We found a total of 90 seedlings from 5 trees and shrubs species in the disturbed plots, while we registered 174 seedlings of 12 species in Old-growth plots. The Standing-dead plots had 83 seedlings from five species, while the Blow-down plots presented 7 seedlings from two species. We did not find any seedling in the total destruction plot. In the Old-growth plots, seedlings were largely dominated by the shade tolerant *A. luma, M. parviflora* and *M. planipes*. In the Standing-dead plots, seedlings were dominated by *C. paniculata* (48% of the total), but also present the shade-tolerant tree *A. luma* (24%), the pioneer tree species *W. trichosperma* and *E. coccineum* (18% and 8% respectively). In the Blow-down plot, *E. coccineum*, a pioneer species, and *A. lanceolata* were the only recorded species.

We found different mechanism of forest regeneration in the plots. In the Old-growth plot most regeneration was from seeds, while in the Standing-dead plots most regeneration become from resprout and few tree seedlings from seeds (Table 4). In the Old-growth plots, most of the tree seedlings corresponded to species with fleshy fruits dispersed by birds, while in the Standing-dead plots most seeds were wind-dispersed. In the Blow-down plots, all seedlings came from seeds (Table 4), but a half of the seedlings corresponded to species dispersed by birds (e.g. *Ribes magellanicum*) and the other half were species dispersed by wind (e.g. *E. coccineum*). Second, in all plots most seedlings were growing on organic substrata (Table 4), such as organic soil on the ground, root discs and logs, while few seedlings were growing directly either on the tephra or on the rocks. In the Blow-down plot we found 7 seedlings of *E. coccineum*, growing just in remnants of organic litter; in Standing-dead plot 11 seedlings growing directly in the tephra, while the other 72 seedlings were resprouting directly from the standing tree’s trunks. In the Old-growth plots we found just one species growing in an exposed rock while the rest of the seedlings were growing in organic substrates (Table 4).

**Table 4.** Number of seedlings per regeneration type and substratum type in the plots.

|  | Regeneration type |  | Substratum type |  |
| --- | --- | --- | --- | --- |
|  | Resprouting | Seed | Mineral | Organic |
| Total-destruction | - | - | - | - |
| Blow-down | - | 7 | - | 7 |
| Standing-dead | 74 | 9 | 11 | 72 |
| Old-growth | 31 | 127 | 1 | 157 |

Other small plant species in the studied plots (Table 1) showed differences along the disturbance gradient (Table 2). For example, the Old-growth plot has species in all categories of plant cover, with few species covering over 75% of the ground, and many species covering between 1-5% (Fig. 6). In contrast, Standing-dead and Blow-down plots were covered by fewer species, and concentrated in the lower ranges of coverage (Fig. 6). In the Total destruction plot we only found a non-vascular plant (*Marchantia* sp.) with a cover between 5-25 % (Fig. 6) growing on a log exposed to the surface. The dominant understory species in the Old-growth forest were the bamboo *Chusquea* spp. and the fern *L. quadripinnata.* These species were also present in the Standing-dead and Blow-down plots, although scattered and with a low percent of coverage in the disturbed plots.

**Figure 6.**
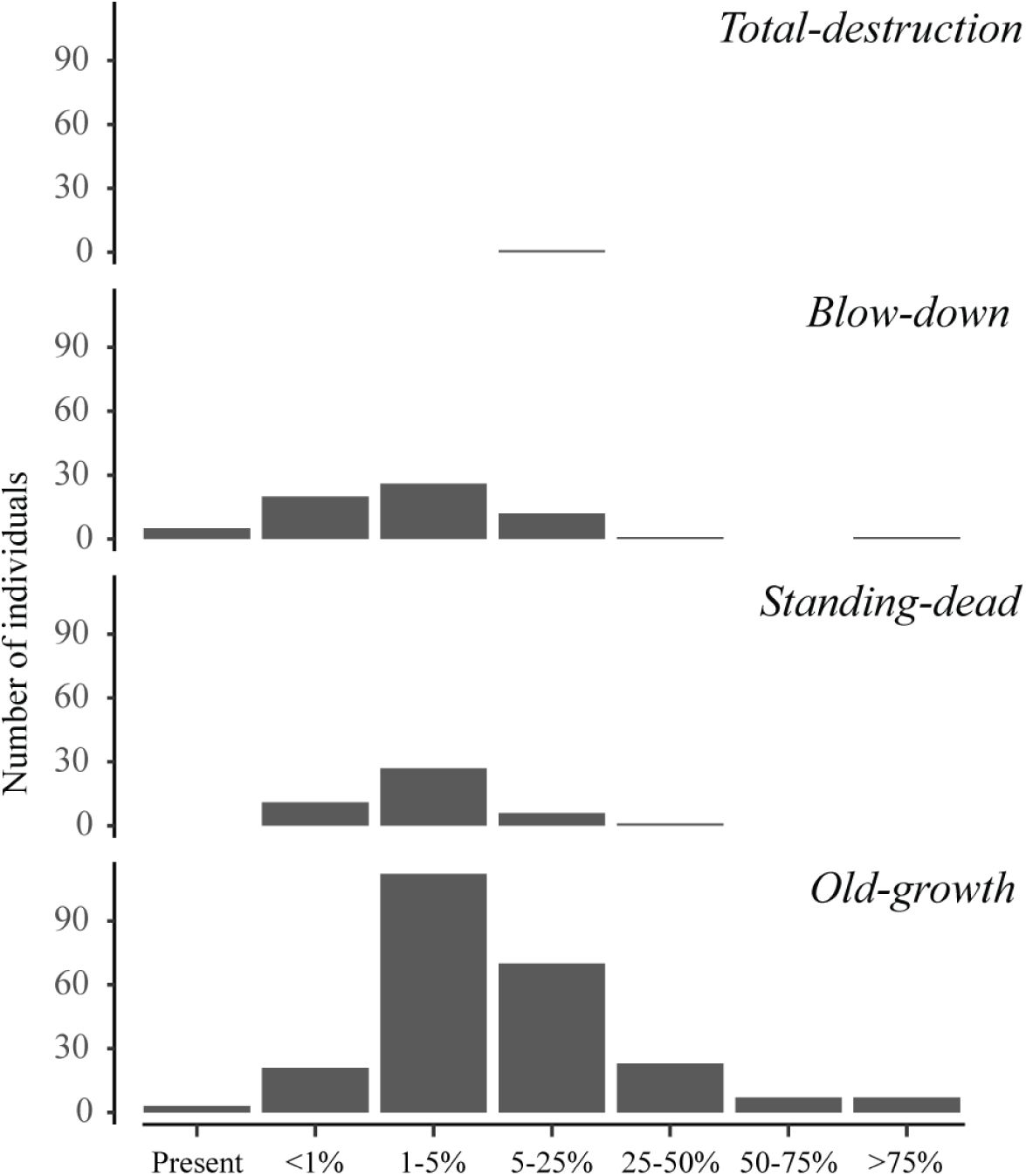
Plant cover (including vascular and non-vascular species) at the ground level of the study areas affected by the Chaitén volcano.

## 4. Discussion

### 4.1 Early vegetation establishment: the relevance of biological legacies

Few studies have been conducted short after an eruption in the southern Andes, however successional model predict that disturbed areas in low and mid altitude (<1000 m a.s.l.) forests would be dominated by *Nothofagus* species due to the shade-intolerance (e.g. Veblen and Ashton, 1978). However, in this study after the first year we did not register *Nothofagus* species despite were abundant in the surroundings non disturbed areas. It might be due to cyclic seed production with not constant seed rain along the years (Donoso, 1993; Veblen, 1985), and the short-term and limited spatial extension we were able to monitor the vegetation. However, our results pointed that regeneration occurs mainly due to an *in situ* development of plants associated to biological legacies such as logs, soil clumps and individuals that survive to the eruption. Other areas disturbed by Chaiten volcano presented similar patterns of types and abundance of biological structures (Swanson et al., 2013).

The initial establishment of the tree species, beside of the disturbance gradient, may be influenced by the interaction between life history of the species and the altitudinal gradient. Life history attributes such as seed dispersal, flowering phenology, growth form, resprouting ability or light-temperature stress resistance are a mechanism that influence the recovery response of the post-disturbance vegetation (Eriksson 2000; McIntyre et al., 1994; McLachlan and Bazely, 2001; Pausas and Lavorel, 2003; Tilman, 1997). In the disturbed plots, the principal mechanism of regeneration was the resprouting capability of some species (Tables 1 and 4). This is a frequent strategy in many plant species in response to disturbances of different types and intensities (Bellingham and Sparrow, 2000; Bond and Midgley, 2001). The seed dispersion syndrome could be also an important mechanism that influence the colonization after disturbance. Chilean forests have one of the highest proportions of plant dispersed by frugivorous birds compared to other temperate rainforests (Armesto et al. 1987, Willson et al., 1996). Birds consume a high proportion of fruits and may disperse the seed to specific places, such as to a perch (Armesto et al., 2001; Lindenmayer and Franklin, 2002). The seed availability might increase in sites where remnant snags, shrubs or perching trees where available (e.g., Bustamante-Sánchez, 2011). Finally, among the wind dispersed seeds, we only found seedlings of *E. coccineum* in disturbed plots. While this tree is adapted to stressful condition, the seed is able to disperse for long-distances (Donoso, 2006).

For non-tree species, Old-growth plot had more species than Standing-dead plot. On the contrary, for tree species Standing-dead plot is richest than Old-growth, revealing the effect of the eruption on the small plant species. On the other hand, in Blow-down and in Total-destruction plot, i.e. from mid to high disturbance intensity at mid to high altitude, and without or sparse biological legacies, we found low species richness and coverage, and sparse tree regeneration. This mosaic of legacies in the landscape offers opportunities for the recolonization of many different species in the affected area. The different species composition would promote different pathways to early succession with different communities, but also different recovery rates. We propose that in low to mid elevation areas, the vegetation recovery will be faster, dominated by species shade-tolerant to semi-tolerant and dispersed by birds, mostly due to biological legacies. On the other hand, at higher altitude, the vegetation recovery would be slower, recolonization made by shade-intolerant and seed dispersed by winds because the most stressful conditions of temperature at higher altitudes.

In synthesis, the results point to an early succession with (i) a mosaic of pioneering-wind dispersed species, scattered survivors regrowing and spreading from biological legacies, and plant species dispersed by frugivorous birds, likely favored by the biological legacies; (ii) the early succession is influenced by the interaction of the species-specific life history (including invasion by exotic species), altitudinal gradient and the different intensity of disturbance in a broad area.

## Acknowledgements

Field work was financed by CONICYT PDA-24. A.L. and D.C. acknowledge support from grant CONICYT/FONDAP/15110009. RM acknowledge the scholarship DAAD/Becas Chile during late stages of this manuscript.

## Notes

#### Summary of Updates

Some grammar mistake have been corrected in the abstract

